# Activation of DMH GABAergic neurons, but not local GABAergic AgRP neurons, attenuates chronic stress-induced POMC neuron hyperactivity

**DOI:** 10.64898/2026.04.01.715870

**Authors:** Yuting Chen, Amin Karimi Moghaddam, Quansheng Du, Yun Lei, Xin-Yun Lu

## Abstract

Identifying the neural circuits engaged and reshaped by chronic stress is critical for understanding how adaptive responses shift to maladaptive behaviors that contribute to stress-related disorders. Our previous work demonstrates that chronic unpredictable stress (CUS) induces a persistent increase in the firing activity of proopiomelanocortin (POMC) neurons in the arcuate nucleus (ARC). This hyperactivity is due, in part, to a reduction in GABAergic synaptic transmission onto POMC neurons, indicating a disruption in inhibitory control. However, the sources of GABAergic inputs responsible for this effect of chronic stress are unknown. Although AgRP neurons provide local GABAergic input onto POMC neurons and are suppressed by chronic stress, chemogenetic activation of AgRP neurons during stress exposure failed to reduce POMC neuron hyperactivity. GABAergic projections originating from the dorsomedial hypothalamus (DMH) represent another source of inhibitory input to POMC neurons. We found that CUS decreased the firing activity of DMH GABAergic neurons with sex differences, with females exhibiting greater vulnerability to stress-induced suppression. Chemogenetic activation of these neurons during chronic stress markedly attenuated POMC neuron hyperactivity in both sexes, indicating that DMH GABAergic neurons function as a critical upstream regulator of POMC neuron activity under chronic stress. These findings suggest that reduced inhibitory input from DMH GABAergic neurons, rather than local GABAergic AgRP neurons, drives POMC neuron hyperactivity. The weakening of the DMH^GABA^→ARC^POMC^ circuit activity may represent a novel mechanism underlying maladaptive stress responses and a potential therapeutic target for stress-related disorders.

## INTRODUCTION

Chronic stress can induce maladaptive responses that impair cognitive, emotional, and social functioning by altering neuronal activity, neural circuitry, and endocrine regulation. Defining how these stress-sensitive networks are remodeled is critical for elucidating the mechanisms that drive the transition from adaptive to maladaptive responses and may reveal potential targets for treating stress-related disorders.

The arcuate nucleus (ARC) is located in the mediobasal hypothalamus, adjacent to the third ventricle, and extends ventrally into the median eminence, allowing circulating hormones to directly influence ARC neurons due to the absence of a typical blood–brain barrier. The ARC contains two distinct populations of neurons that express proopiomelanocortin (POMC) and agouti-related peptide (AgRP) [1, 2]. Accumulating evidence indicates that these neurons are sensitive to psychological stress, integrating hormonal and neural signals that in turn regulate behavioral responses to stress [3–6]. We have previously shown that POMC neurons, to a much less extent AgRP neurons, can be activated rapidly by acute restraint and forced swim stress [3]. However, the firing activity of these neurons responds to chronic stress in opposite directions [5–7]. Chronic unpredictable stress (CUS), a paradigm that generates behavioral deficits such as anhedonia and behavioral despair [5, 6, 8, 9], suppresses AgRP neuron activity while increasing POMC neuron activity [5, 6]. Furthermore, direct stimulation of AgRP neurons can reverse behavioral deficits induced by CUS [5]. By contrast, chronic activation of POMC neurons can mimic the effects of CUS by inducing similar behavioral deficits [6]. Conversely, inhibition of POMC neurons reverses chronic stress-induced behavioral changes [6]. These findings suggest that POMC neuron activity is both necessary and sufficient for mediating chronic stress–induced behavioral phenotypes. This causal role positions POMC neurons as a central node that integrates stress-related signals to drive behavioral adaptations. Further analyses of synaptic transmission in POMC neurons revealed that, while glutamatergic excitatory transmission remained unchanged, GABAergic inhibitory transmission was reduced [6]. This shift in the excitatory–inhibitory balance likely enhances the overall excitability of POMC neurons. Notably, the decreased frequency of spontaneous inhibitory postsynaptic currents (IPSCs) in POMC neurons points to a presynaptic mechanism, implicating circuit-level disinhibition in driving POMC neuron hyperactivity [6]. However, the specific sources of the GABAergic inputs to POMC neurons that are suppressed by CUS have yet to be determined.

One source of GABAergic input to POMC neurons arises locally from AgRP neurons in the ARC. Studies have shown that AgRP neurons project to and synapse onto POMC neurons, providing direct GABAergic inhibition of these cells [1, 10–12]. Supporting this, ablation of AgRP neurons reduces baseline GABAergic tone onto POMC neurons [13], and that optogenetic stimulation of AgRP neurons triggers GABA release onto POMC cells and suppresses their firing, demonstrating a direct, functional synaptic connection between AgRP and POMC neurons [10, 11, 14]. Given that chronic stress inhibits AgRP neuron firing [5], one would expect that this reduces GABAergic input to POMC neurons, thereby contributing to the disinhibition of POMC neuron activity; however, this remains to be determined.

In addition to local inputs, the dorsomedial hypothalamus (DMH) represents a major source of GABAergic input to POMC neurons [15, 16]. The DMH contains both glutamatergic and GABAergic but predominantly GABAergic neurons [17, 18] that project to the ARC [16], providing inhibitory control over POMC neuron activity [15, 16]. DMH neurons have been reported to be activated by acute stressors [19–21], as well as by repeated restraint or chronic variable stress [22], as indicated by c-Fos and FosB expression. However, these markers are used for mapping transcriptionally responsive neurons rather than real-time neuronal firing. With typically low basal expression, they detect neurons activated above a threshold but provide limited information about inhibitory activity. Thus, it remains unclear how chronic stress influences the activity of GABAergic neurons within the DMH, and whether these neurons contribute to reduced GABAergic input and POMC neuron hyperactivity.

In this study, we aimed to determine the contributions of two inhibitory pathways, local GABA-releasing AgRP neurons and DMH GABAergic neurons, to chronic stress–induced POMC neuron hyperactivity. We first tested the hypothesis that reduced AgRP activity diminishes GABA release, leading to disinhibition of POMC neurons, by examining whether chemogenetic activation of AgRP neurons during chronic stress could counteract POMC neuron hyperactivity. Next, because the impact of chronic stress on DMH GABAergic neuron firing is unknown, we assessed how stress alters their firing properties and whether activation of these neurons modulates POMC activity under chronic stress.

## MATERIALS AND METHODS

### Animals

Pomc-GFP mice (Stock No. 009593), AgRP-IRES-Cre knock-in mice (Stock No. 012899) and VGAT-IRES-Cre knock-in mice (Stock No. 028862) were purchased from Jackson Laboratory. In Pomc-GFP mice, GFP expression is driven by the mouse Pomc promoter/enhancer regions [12], enabling visualization of POMC neurons. AgRP-IRES-Cre mice express Cre recombinase under control of the agouti-related peptide (AgRP) locus, selectively targeting AgRP neurons [23]. VGAT-IRES-Cre mice express Cre recombinase selectively in inhibitory GABAergic neurons without disrupting endogenous vesicular inhibitory amino acid transporter expression [17]. Male AgRP-IRES-Cre or VGAT-IRES-Cre mice were crossed with female Pomc-GFP mice to generate Pomc-GFP;AgRP-IRES-Cre and Pomc-GFP;VGAT-IRES-Cre offspring. These mice exhibited GFP fluorescence labeling POMC neurons and Cre expression in AgRP or GABAergic neurons, enabling Cre-dependent manipulation of neuronal activity. Mice were housed in groups of 3-5 mice per individually ventilated cage, with ad libitum access to food and water under a 12-h light-dark cycle (lights on at 06:00 AM). All animal procedures were conducted in accordance with the National Institutes of Health (NIH) Guide for the Care and Use of Laboratory Animals and approved by the Institutional Animal Care and Use Committee of Augusta University.

### Stereotaxic surgery and AAV injection

Mice at 7 weeks of age received stereotaxic microinjections of AAV vectors under anesthesia, as previously described [5, 6, 24]. A Synapsin-driven, Cre-dependent mCherry control virus (AAV5-hSyn-DIO-mCherry; Addgene #50459; hereafter referred to as AAV-DIO-mCherry) or a Cre-dependent hM3D(Gq) DREADD receptor with an mCherry reporter for CNO-induced neuronal activation (AAV5-hSyn-DIO-hM3D(Gq)-mCherry; Addgene #44361; hereafter referred to as AAV-DIO-hM3Dq-mCherry) was injected into the ARC (coordinates: AP −1.4 mm, ML ±0.2 mm, DV −5.8 mm from bregma) of Pomc-GFP;AgRP-IRES-Cre mice, or into the DMH (coordinates: AP −1.6 mm, ML ±0.4 mm, DV −5.4 mm from bregma) of VGAT-IRES-Cre mice and Pomc-GFP;VGAT-IRES-Cre mice. A total volume of 200 nL AAV was delivered at a rate of 40 nL/min using a 30-gauge stainless steel syringe (Hamilton Company, Reno, NV, USA) connected to a UMP3 microsyringe pump (World Precision Instruments, Sarasota, FL, USA). After injection, the syringe was left in place for an additional 5 minutes to allow diffusion and minimize backflow.

For experiments assessing connectivity between AgRP neurons in the ARC or GABAergic neurons in the DMH with POMC neurons, AAV-DIO-hM3Dq-mCherry was injected unilaterally into the ARC of Pomc-GFP;AgRP-IRES-Cre mice or into the DMH of Pomc-GFP;VGAT-IRES-Cre mice.

### Fluorescent imaging

Mice were transcardially perfused with 0.1 M phosphate-buffered saline (PBS) followed by 4% paraformaldehyde (PFA) in PBS. Brains were post-fixed overnight in 4% PFA and then cryoprotected in 30% sucrose. Coronal brain sections (40 µm) were cut using a cryostat and stored in cryoprotectant solution. Free-floating sections were incubated with 4’,6-diamidino-2-phenylindole (DAPI; Thermo Fisher Scientific, Waltham, MA, USA) for nuclear labeling and mounted onto glass slides. Fluorescent images were acquired using a Keyence microscope (Keyence, Itasca, IL, USA) with a 4× objective and a Leica confocal microscope (Leica Microsystems, Deerfield, IL, USA) with a 20× objective.

### Chronic unpredictable stress (CUS) procedure and chemogenetic manipulation

Fourteen days after AAV injection, mice were subjected to the CUS protocol as described in our previous studies [5, 6, 8, 9, 25]. Mice were individually housed and exposed to a variety of stressors at unpredictable times of day for 10 consecutive days. Stressors included 2-h restraint, 15-min tail pinch, 24-h constant light, 24-h wet bedding with a 45° cage tilt, 10-min scrambled inescapable foot shocks, and 30-min elevated platform exposure. All stress procedures were conducted in a dedicated procedure room, whereas control mice were group-housed and received brief daily handling in the housing room.

For chemogenetic neuronal activation, mice were administered clozapine-N-oxide (CNO; Sigma-Aldrich, St. Louis, MO, USA) in drinking water at 5 mg/L, beginning one day prior to CUS and maintained throughout the 10-day stress period. After the final stressor, CNO-containing water was replaced with regular drinking water.

### Whole-cell current-clamp recordings

Electrophysiological recordings were performed one day after the completion of the 10-day unpredictable stress paradigm to avoid acute stress effects and assess POMC neuron firing as a measure of chronic stress-induced drive as previously described [5, 6]. Control and CUS mice were anesthetized with isoflurane before decapitation. Brains were quickly dissected in an ice-cold oxygenated (95% O_2_/5% CO_2_) cutting solution (210 mM sucrose, 3 mM KCl, 2 mM MgCl_2_, 2 mM CaCl_2_, 1.25 mM NaH_2_PO_4_, 10 mM D-glucose, and 24 mM NaHCO_3_, pH 7.4 and osmolarity of 300 mOsm/L). Coronal brain slices at 300 μm thickness were prepared with a Leica VT1200S Vibratome (Leica Microsystems, Deerfield, IL, USA). Slices were then transferred to an oxygenated artificial cerebrospinal fluid solution (aCSF; 124 mM NaCl, 2 mM KCl, 2 mM MgSO_4_, 2 mM CaCl_2_, 1.25 mM NaH_2_PO_4_, 26 mM NaHCO_3_, and 10 mM Glucose, pH 7.4 and osmolarity of 300 mOsm/L) at 32 °C for 60 min and subsequently at room temperature for another 30 min prior to recordings. Slices were transferred to the recording chamber and continuously perfused with oxygenated aCSF at a flow rate of 1-2 ml/min at room temperature.

Neurons were visualized with a fixed stage upright microscope Examiner.A1 (Zeiss, White Plains, NY, USA) using 5× objective and 40× water-immersion objective with infrared differential interference contrast optics (IR-DIC) and fluorescence optics. POMC neurons were identified based on their anatomical location within the ARC of the hypothalamus and by GFP fluorescence, visualized by Calibri.2 illumination system (Zeiss) in combination with a filter. DMH GABAergic neurons were identified by location in the dorsomedial hypothalamus and by mCherry fluorescence. Patch electrodes were prepared by a P-1000 micropipette puller (Sutter Instrument, Novato, CA, USA). Pipettes were filled a potassium gluconate-based internal solution (120 mM potassium gluconate, 20 mM KCl, 2 mM MgCl_2_, 10 mM HEPES, 2 mM ATP, 0.25 mM GTP, and 0.1 mM EGTA, pH 7.4 and osmolarity of 295 mOsm/L) with a final resistance of 3–5 MΩ. All recordings were made using a MultiClamp 700B Microelectrode Amplifier (Molecular Devices, San Jose, CA, USA), and data was filtered at 2 kHz and digitized at 10 kHz by using Axon Digidata 1550A (Molecular Devices) and analyzed on a PC computer with pCLAMP 10.7 program (Molecular Devices). The series resistance was measured before and after patch clamp recording. Only those cells with series resistance less than 20 MΩ and varied less than 15% during the recording were included for analysis. Membrane potential and spontaneous action potential (AP) firing rates were measured by whole-cell patch-clamp recording in current-clamp mode. Neurons were classified as quiet or active based on firing rates: neurons with firing rates **<** 0.5 Hz were considered quiet, whereas those with firing rates ≥ 0.5 Hz were considered active. The interspike intervals (ISIs) for each neuron were measured; coefficient of variations (the ratio of the standard deviation of ISI to the mean of ISI) were calculated. The AP threshold was defined as membrane potential at which the first derivative of the voltage (dV/dT) exceeded 10 mV/ms. AP amplitude, half width, rise time (10%-90%), decay time (10-90%) and afterhyperpolarization (AHP) amplitude were determined by pCLAMP using AP threshold as baseline. And the peak-to-AHP time was defined as the interval between AP peak and AHP peak.

### Statistical analysis

All results were presented as mean ± s.e.m. (standard error of the mean). Statistical analyses were performed with GraphPad Prism 10 (GraphPad Software, Inc., CA, USA). For normally distributed data, two-tailed t test was used to assess differences between two experimental groups with equal variances, while unpaired t test with Welch’s correction was used for comparisons between two groups with unequal variances. For normal distributed data involving more than two groups, one-way analyses of variance (ANOVAs) followed by Bonferroni post hoc tests were used. For non-normally distributed data, Mann-Whitney U test was performed to compare two groups and Kruskal-Wallis test followed by Dunn’s multiple comparisons was used for three or more groups. P < 0.05 was considered statistically significant.

## RESULTS

### Chronic activation of AgRP neurons during chronic stress fails to reduce POMC neuron hyperactivity

Previous work has shown that chronic exposure to unpredictable stress (CUS) produces a persistent decrease in AgRP neuron firing, measured one day after the last stressor to avoid acute stress effects [5]. This reduction in AgRP neuron activity may lead to decreased GABA release onto POMC neurons, thereby contributing to their reduced inhibitory tone. If this is the case, restoring AgRP neuron activity would be expected to reverse POMC neuron disinhibition. To test this hypothesis, we examined whether chemogenetic stimulation of AgRP neurons during stress exposure could attenuate CUS-induced POMC neuron hyperactivity. This approach allows for a direct assessment of the functional role of AgRP neuron–mediated GABAergic signaling in regulating POMC neuron activity under chronic stress conditions.

To enable chemogenetic activation of AgRP neurons and assess their effects on POMC neurons, Pomc-GFP mice were crossed with AgRP-IRES-Cre mice to generate Pomc-GFP;AgRP-IRES-Cre mice (Fig. 1A). To verify the efficiency of Cre-dependent mCherry labeling in AgRP neurons and their terminals, as well as their anatomical relationship with GFP-labeled POMC neurons, mice received unilateral injections of an AAV expressing Cre-dependent mCherry (Fig. 1B-1D). Fluorescent imaging revealed mCherry-positive AgRP terminals closely apposed to GFP-positive POMC neurons (Fig. 1E), consistent with the formation of synaptic contacts between these two neuronal populations [12].

**Figure 1.**
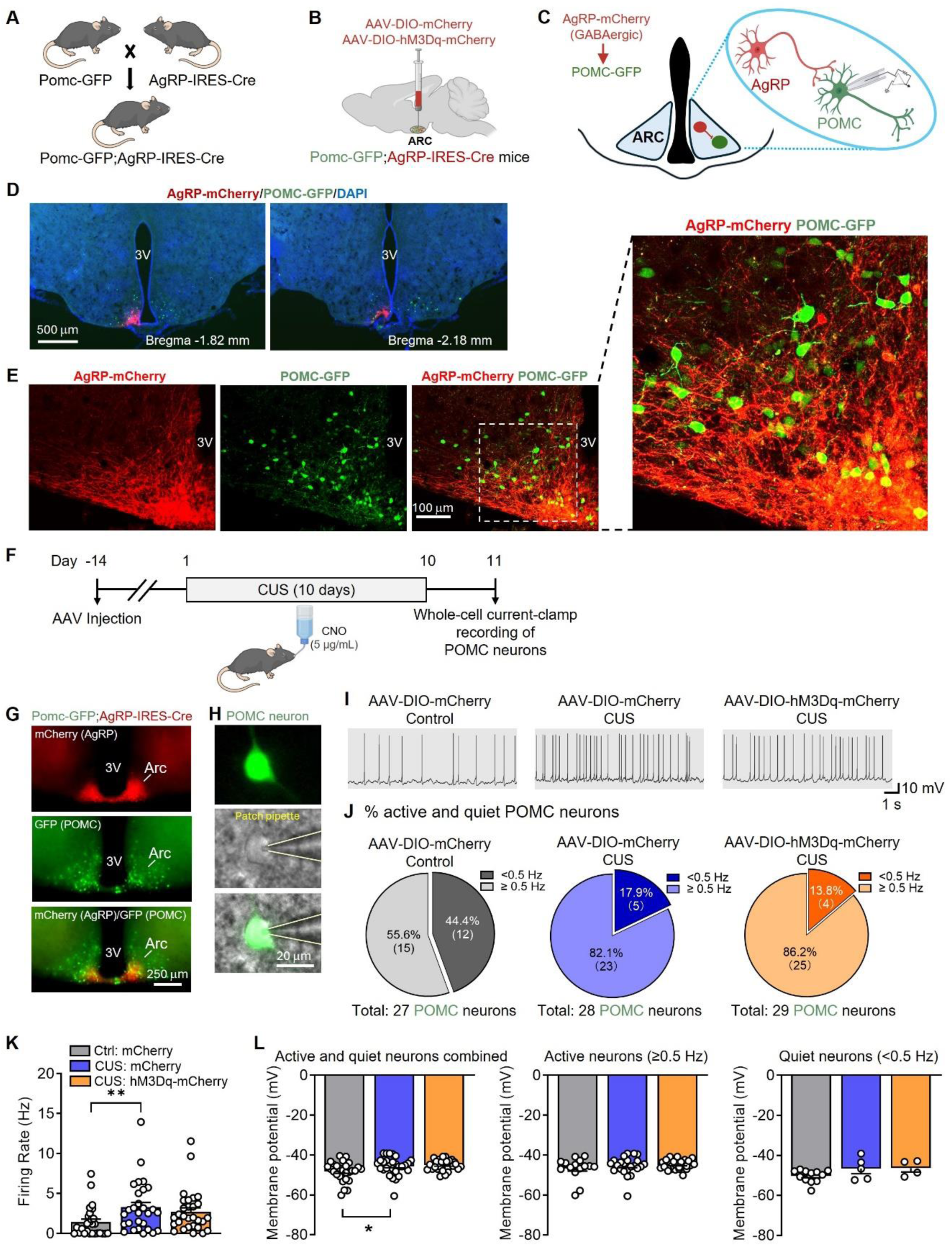
Chronic chemogenetic activation of AgRP neurons during chronic stress exposure has no effect on CUS-induced POMC neuron hyperactivity. **A** Breeding strategy to generate *Pomc-GFP;AgRP-IRES-Cre* mice. **B** Illustration of AAV injection in the arcuate nucleus (ARC). **C** Schematic illustrating the projections from AgRP neurons to POMC neurons in the ARC. **D** Unilateral intra-ARC injection of AAV-DIO-hM3Dq-mCherry in *Pomc-GFP;AgRP-IRES-Cre* mice. Red, mCherry-labeled AgRP neurons and terminals. Green, GFP-labeled POMC neurons and processes. **E** Representative images showing AgRP terminals (red) in close apposition to POMC neurons (green) in the ARC. **F** Experimental timeline of intra-ARC AAV injection, chronic stress procedure with administration of CNO, and whole-cell current-clamp recording of POMC neurons. **G** Representative images depicting mCherry-labeled AgRP neurons and GFP-labeled POMC neurons in the ARC of mice that received bilateral intra-ARC AAV injections. **H** Representative images showing whole-cell patch-clamp recording of POMC neurons. **I** Representative traces of action potentials from three treatment groups. **J** The percentage of active and quiet POMC neurons and total number of neurons recorded. **K** Spontaneous firing rate of POMC neurons (Kruskal-Wallis test, *P* = 0.0055). **L** Membrane potential of POMC neurons. Left: all active and quiet neurons combined (Kruskal-Wallis test, *P* = 0.0174); middle: active neurons only (Kruskal-Wallis test, *P* = 0.8542); right: quiet neurons only (*F*_(2,18)_ = 2.419, *P* = 0.1174). Control (Ctrl)-AAV-DIO-mCherry group (Ctrl: mCherry): total 27 neurons from 3 male mice. Chronic unpredictable stress (CUS)-AAV-DIO-mCherry group (CUS: mCherry): total 28 neurons from 3 male mice; CUS-AAV-DIO-hM3Dq-mCherry (CUS: hM3Dq-mCherry): total 29 neurons from 3 male mice. \**P* < 0.05, \*\**P* < 0.01 compared with the CUS: mCherry group.

To induce sustained stimulation of AgRP neurons during chronic stress, Pomc-EGFP;AgRP-IRES-Cre mice were injected in the ARC with AAVs expressing either Cre-dependent control mCherry or hM3Dq-mCherry for chemogenetic activation. To activate hM3Dq-expressing AgRP neurons, our pilot studies initially delivered CNO via daily intraperitoneal (i.p.) injections over the 10-day stress exposure period, with control mice receiving the same treatment. One day after the final stressor, whole-cell current-clamp recordings of GFP-labeled POMC neurons were performed to assess the effects of chronic AgRP neuron activation on spontaneous firing activity of POMC neurons. No differences in POMC neuron firing were observed between CUS mice expressing hM3Dq and those expressing control mCherry (data not shown). It appeared that repeated i.p. injections themselves induced stress-related effects in control mice, creating a confounding variable that complicated interpretation. To avoid the stress-related confounds associated with repeated intraperitoneal injections, CNO was subsequently administered in the drinking water throughout the 10-day stress paradigm (Fig. 1F). Whole-cell current-clamp recordings of POMC neurons were then performed as described above (Fig. 1G–I). Consistent with our previous findings [6], CUS increased the proportion of active POMC neurons, reflected by a reduction in the percentage of “quiet” neurons (defined by firing rates < 0.5 Hz) (Fig. 1J), elevated the overall firing rates and depolarized their membrane potential (Fig. 1K, 1L). However, chronic chemogenetic activation of AgRP neurons via hM3Dq with CNO in drinking water did not significantly alter any of these measures (Fig. 1J-1L). These findings suggest that increasing AgRP neuron activity alone is insufficient to restore inhibitory control over POMC neurons under chronic stress conditions.

### Chronic stress decreases spontaneous firing of GABAergic neurons in the DMH

GABAergic neurons in the DMH have been identified as an additional source of inhibitory input to POMC neurons in the ARC [15]. If these neurons contribute to the CUS-induced reduction in inhibitory tone onto POMC neurons, their activity would be expected to decrease under chronic stress. To test this, AAVs expressing Cre-dependent mCherry were injected into the DMH of VGAT-IRES-Cre mice to selectively label GABAergic neurons for electrophysiological recordings. Whole-cell current-clamp recordings were performed one day after the 10-day CUS paradigm (Fig. 2A–2D) to assess spontaneous firing activity. Data from male and female mice were initially pooled and then analyzed separately to evaluate potential sex differences in the effects of chronic stress on DMH GABAergic neuron activity. We found that the percentages of active GABAergic neurons were comparable between sexes under basal conditions (male: 45.5% vs. female: 48.5%). Following CUS exposure, the percentage of active neurons was decreased similarly in both sexes (male: 19.0% vs. female: 17.2%) (Fig. 2E, 2H and 2K). However, spontaneous firing rates differed significantly between sexes. Under basal conditions, female mice exhibited much higher firing rates than male mice (male: 0.5223 ± 0.1242 Hz; female: 1.193 ± 0.2776 Hz) (Fig. 2F, 2I and 2L). While CUS reduced firing rates in both sexes (male: 0.3457 ± 0.1686 Hz; female: 0.280 ± 0.0878 Hz), the magnitude of reduction was greater in females (∼76.5%) than in males (∼33.8%), suggesting that DMH GABAergic neurons in females are more sensitive to chronic stress–induced suppression. To further characterize firing patterns, we analyzed interspike interval distributions and the coefficient of variation (CV) as measures of spike regularity. CUS did not alter the cumulative frequency distribution of interspike intervals or the CV, whether data were pooled or analyzed by sex (Fig. 2G, 2J, 2M). However, correlation analysis revealed a significant negative relationship between firing rate and CV in control mice, which was absent in CUS mice (Fig. 2G, 2J, 2M). These results indicate that CUS suppresses spontaneous firing activity in DMH GABAergic neurons without changing overall spike timing variability, but disrupts the normal relationship between firing rate and firing regularity.

**Figure 2.**
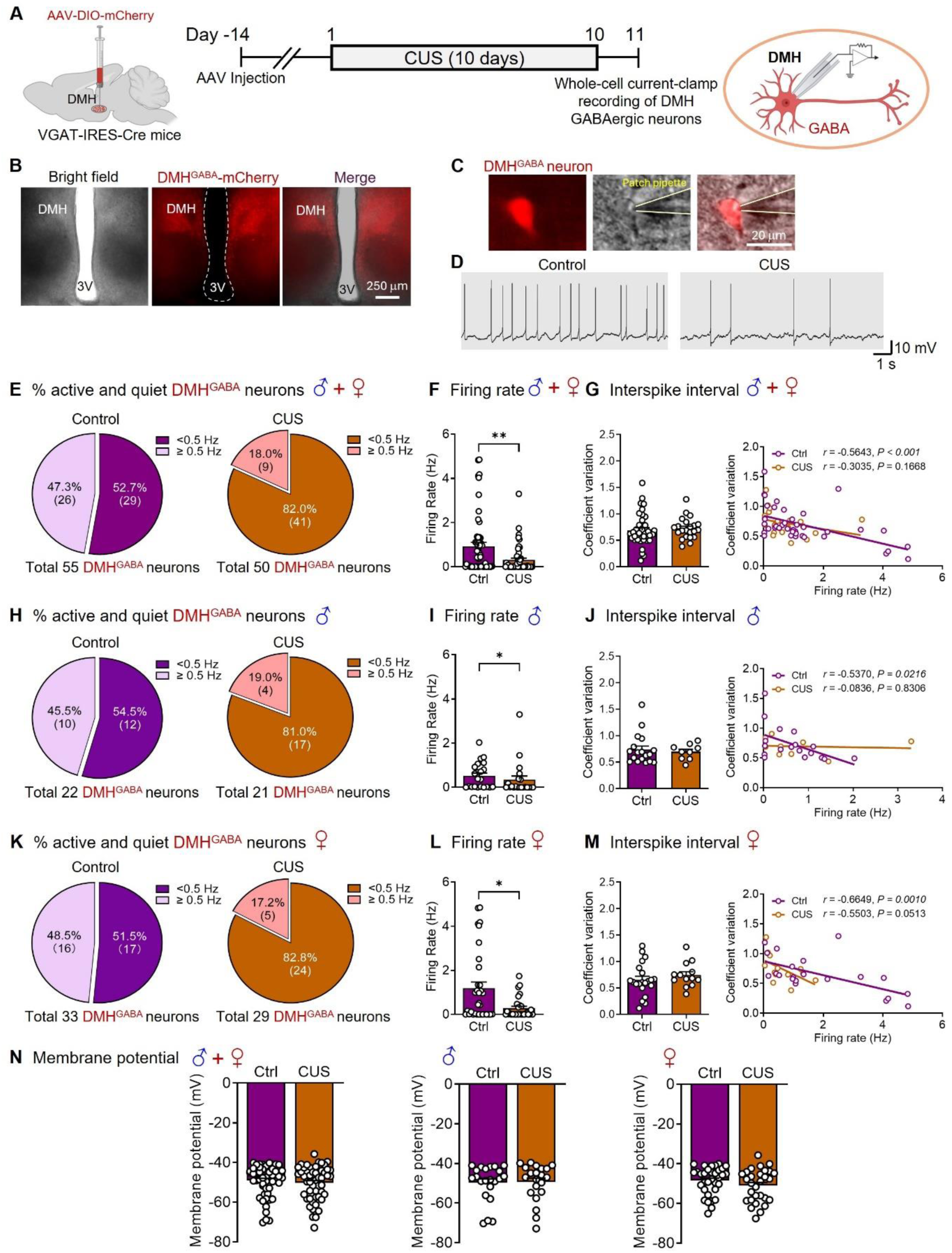
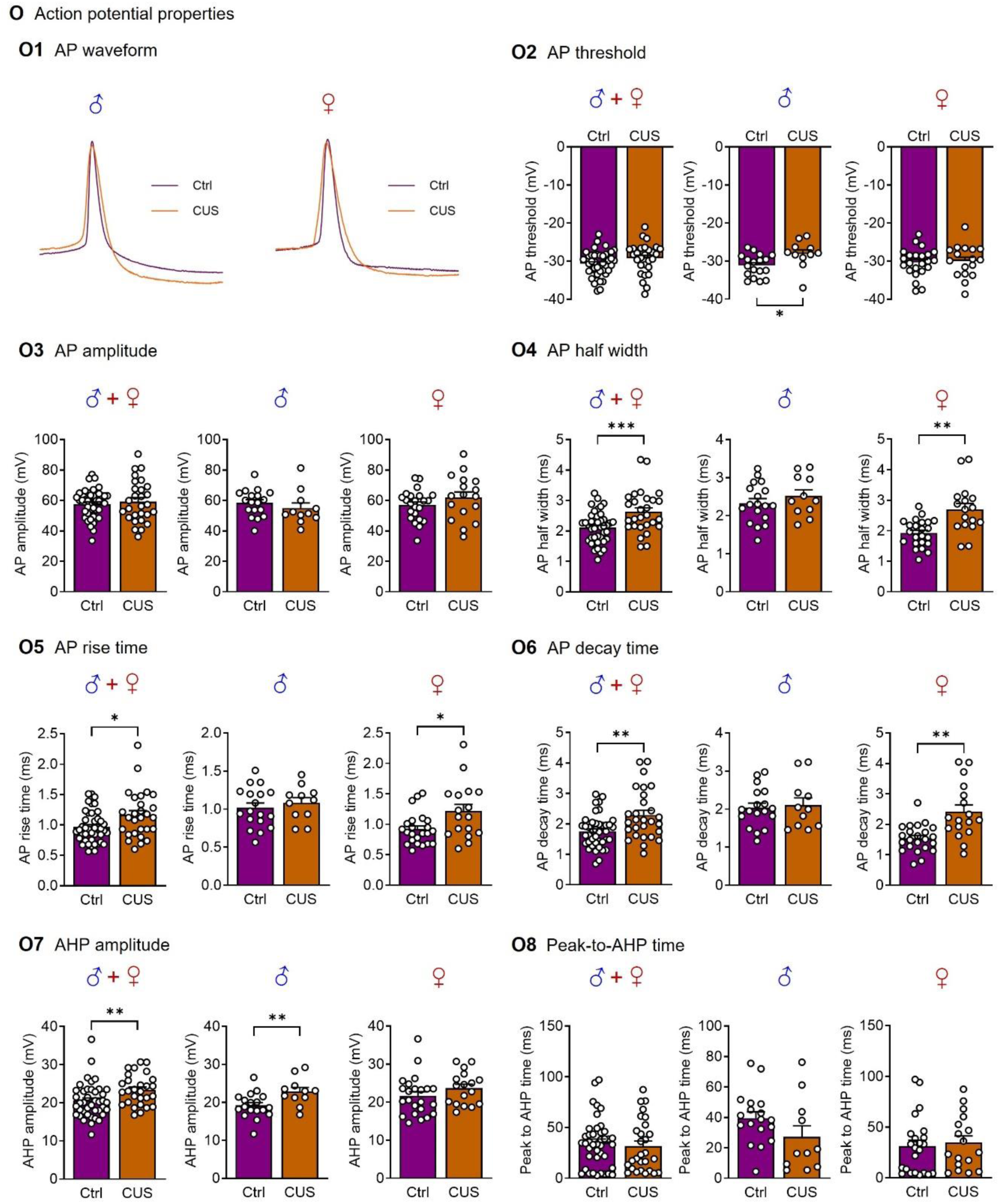
CUS decreases spontaneous firing rates of GABAergic neurons in the DMH. **A** Timeline of experimental procedures. VGAT-IRES-Cre mice received bilateral intra-DMH injection of AAV-DIO-mCherry to label GABAergic neurons. **B** Representative bright-field and fluorescence images showing mCherry-labeled GABAergic neurons within the DMH. **C** Representative images showing whole-cell patch-clamp recording of DMH GABAergic neurons. **D** Representative traces of action potentials from control and CUS mice. The percentages of active and quiet DMH GABAergic for combined male and female mice (E), males only (H), and females only (K). Firing rates of DMH GABAergic neurons for combined male and female mice (F), males only (I), and females only (L) (Combined: Mann Whitney test, *P* = 0.0022; Male: Mann Whitney test, *P* = 0.0451; Female: Mann Whitney test, *P* = 0.0216). Firing patterns (mean coefficients of variation) and the correlation between firing rate and coefficient of variation of DMH GABAergic neurons recorded from combined male and female mice (G), males only (J), and females only (M) (Average coefficients of variation: Combined: Mann Whitney test, *P* = 0.2786; Male: Mann Whitney test, *P* = 0.7814; Female: Mann Whitney test, *P* = 0.2182). **N** Membrane potential. Left, combined male and female mice (Mann Whitney test, *P* = 0.4269); middle, males only (Mann Whitney test, *P* = 0.7409); right, females only (Mann Whitney test, *P* = 0.1921). **O** Action potential (AP) properties. O1. Representative AP waveforms recorded from DMH GABAergic neurons of control and CUS mice. O2. AP threshold. Left: combined male and female mice (t_(67)_ = 1.592, *P* = 0.1160); middle: males (t_(27)_ = 2.444, *P* = 0.0214); right: females (t_(38)_ = 0.2779, *P* = 0.7826). O3. AP amplitude. Left: combined male and female mice (Unpaired t test with Welch’s correction, *P* = 0.5911); middle: males (t_(27)_ = 0.9406, *P* = 0.3552); right: females (t_(38)_ = 1.287, *P* = 0.2060). O4. AP half width. Left: combined male and female mice (t_(67)_ = 3.613, *P* < 0.001); middle: males (t_(27)_ = 0.9590, *P* = 0.3460); right: females (Unpaired t test with Welch’s correction, *P* = 0.0014). O5. AP rise time. Left: combined male and female mice (Mann-Whitney test, *P* = 0.0252); middle: males (t_(27)_ = 0.6624, *P* = 0.5133); right: females (Mann-Whitney test, *P* = 0.0223). O6. AP decay time. Left: combined male and female mice (Unpaired t test with Welch’s correction, *P* = 0.0038); middle: males (t_(27)_ = 0.3093, *P* = 0.7595); right: females (Unpaired t test with Welch’s correction, *P* = 0.0014). O7. AHP amplitude. Left: combined male and female mice (Mann-Whitney test, *P* = 0.0058); middle: males, (t_(27)_ = 2.798, *P* = 0.0094); right: females (t_(38)_ = 1.313, *P* = 0.1972). O8. Peak-to-AHP time. Left: combined male and female mice (Mann-Whitney test, *P* = 0.7044); middle: males (Mann-Whitney test, *P* = 0.0922); right: females (Mann-Whitney test, *P* = 0.4855). Control (Ctrl): n=55 neurons from 3 male (22 neurons) and 4 female (33 neurons) mice; chronical unpredictable stress (CUS): n=50 neurons from 3 male (21 neurons) and 4 female (29 neurons) mice. \**P* < 0.05, \*\**P* < 0.01, \*\**P* < 0.001 compared with the control group.

In contrast, resting membrane potential did not differ significantly between sexes under either basal or chronic stress conditions (Fig. 2N). Control female and male mice exhibited comparable membrane potentials (male: −49.72 ± 1.967 mV; female: −48.46 ± 1.217 mV), as did CUS-exposed mice (male: −49.41 ± 2.036 mV; female: −51.01 ± 1.524 mV). To further explore the mechanisms underlying sex-specific differences in spontaneous firing rates, we analyzed action potential (AP) waveform properties, including AP threshold, amplitude, half-width, rise time, decay time, afterhyperpolarization (AHP) amplitude and peak-to-AHP time (the interval between the peak of AP and the peak of AHP) (Fig. 2O). These parameters reflect intrinsic membrane excitability and ion channel function that govern spike initiation and repolarization. We observed sex-specific differences in AP waveform properties following CUS. In male mice, CUS increased AP threshold and AHP amplitude (Fig. 2O2, 2O7), whereas in female mice, CUS increased AP half-width (Fig. 2O4), rise time (Fig. 2O5), and decay time (Fig. 2O6). AP amplitude (Fig. 2O3) and peak-to-AHP time (Fig. 2O8) were unaffected in both sexes. These data suggest that chronic stress induces sex-specific adaptations in the kinetics of DMH GABAergic neuron action potentials, which may contribute to the observed differences in firing properties between male and female mice.

### Chronic activation of DMH GABAergic neurons during chronic stress attenuates POMC neuron hyperactivity

Since DMH GABAergic neurons project to ARC POMC neurons [16], CUS-induced inhibition of these neurons may lead to disinhibition of POMC neurons, providing a potential synaptic mechanism for stress-induced POMC hyperactivity. To determine whether activation of DMH GABAergic neurons during chronic stress could counteract this hyperactivity, we crossed Pomc-GFP mice with VGAT-IRES-Cre mice to generate Pomc-GFP;VGAT-IRES-Cre mice (Fig. 3A).

**Figure 3.**
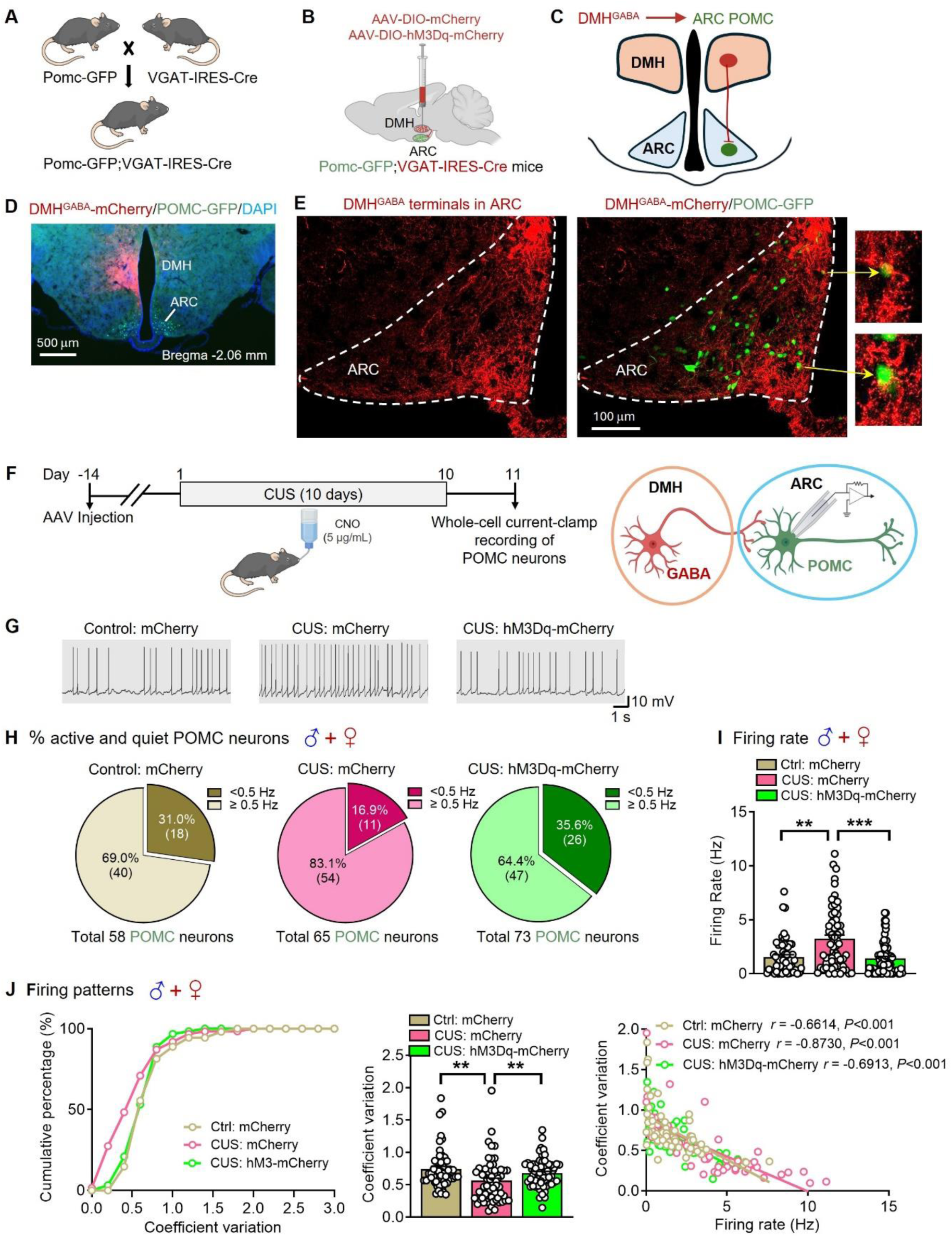
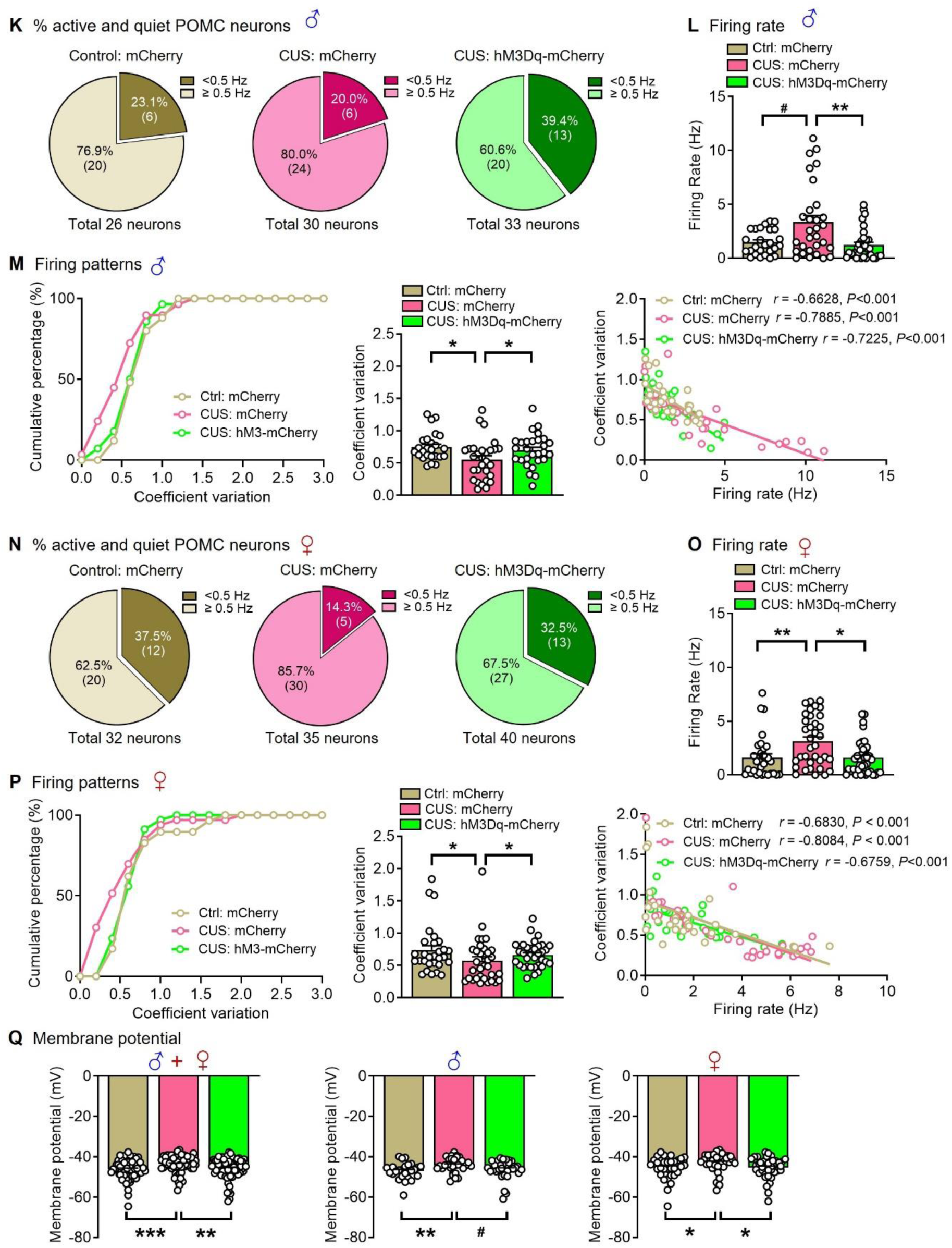
Chronic chemogenetic activation of DMH GABAergic neurons during chronic stress exposure attenuates CUS-induced POMC neuron hyperactivity. **A** Breeding strategy for generating *Pomc-GFP;VGAT-IRES-Cre* mice. **B** Illustration of AAV injection into the DMH. **C** Schematic showing projections from DMH GABAergic neurons to ARC POMC neurons. **D** Unilateral intra-DMH injection of AAV-DIO-hM3Dq-mCherry in *Pomc-GFP;VGAT-IRES-Cre* mice. Red, mCherry-labeled GABAergic neurons and terminals originating from the DMH. Green, GFP-labeled POMC neurons and their processes. **E** Representative images showing GABAergic terminals from the DMH (red) in close apposition to POMC neurons (green) in the ARC. **F** Experimental timeline depicting intra-DMH AAV injections, chronic stress exposure with CNO administration, and whole-cell current-clamp recording of POMC neurons. **G** Representative traces of action potentials from three treatment groups. Percentages of active and quiet POMC neurons from combined male and female mice (H), males only (K), and females only (N). Firing rates of POMC neurons in combined male and female mice (I; Kruskal-Wallis test, *P* < 0.001), males only (L; Kruskal-Wallis test, *P* = 0.0117(^#^*P* = 0.0736)), and females only (O; Kruskal-Wallis test, *P* = 0.0060). Firing patterns (mean coefficients of variation): left, cumulative probability distributions of coefficients of variation. Middle, average coefficients of variation (Combined: Kruskal-Wallis test, *P* = 0.0016; Males: Kruskal-Wallis test, *P* = 0.0198; Females: Kruskal-Wallis test, *P* = 0.0576). Right, correlation analysis between firing rates and coefficients of variation.of POMC neurons recorded from combined male and female mice (J), males only (M), and females only (P). **Q** Membrane potential. Left: combined male and female mice (Kruskal-Wallis test, *P* < 0.001); middle: males only (Kruskal-Wallis test, *P* = 0.0125 (^#^*P* = 0.0581)); right: females only (Kruskal-Wallis test, *P* = 0.0207). Control (Ctrl)-AAV-DIO-mCherry group (Ctrl: mCherry): total 58 neurons from 3 male (26 neurons) and 4 female (32 neurons) mice. Chronic unpredictable stress (CUS)-AAV-DIO-mCherry group (CUS: mCherry): total 65 neurons from 4 male (30 neurons) and 5 female (35 neurons) mice; CUS-AAV-DIO-hM3Dq-mCherry (CUS: hM3Dq-mCherry): total 73 neurons from 4 male (33 neurons) and 5 female (40 neurons) mice. \**P* < 0.05, \*\**P* < 0.01, \*\*\**P* < 0.001 compared with the CUS: mCherry group. DMH, dorsomedial hypothalamus; ARC, arcuate nucleus.

To verify the anatomical relationship between DMH GABAergic terminals and GFP-labeled POMC neurons, these mice received unilateral injections of an AAV expressing Cre-dependent mCherry into the DMH (Fig. 3B-3D). Confocal imaging revealed that mCherry-positive DMH GABAergic terminals were closely apposed to GFP-labeled POMC neurons in the ARC, consistent with the presence of putative synaptic contacts (Fig. 3E). These observations provide anatomical evidence that DMH GABAergic neurons are positioned to directly inhibit POMC neuron activity. As expected, CUS increased the proportion of active POMC neurons in both male and female mice.

Under basal conditions, female mice had a lower percentage of active POMC neurons (62.5%) compared to male mice (76.9%), however, the proportion of active POMC neurons became higher in females (85.7%) than in males (80%) following CUS (Fig. 3H, 3K and 3N). These results indicate that POMC neurons in females are more strongly activated by chronic stress than those in males.

In addition to changes in the proportion of active neurons, CUS increased the spontaneous firing rates of POMC neurons and reduced the irregularity of their firing (Fig. 3I, 3J, 3L, 3M, 3O and 3P), indicating that chronic stress not only enhances POMC neuron activity but also stabilizes their firing patterns. These CUS-induced effects were attenuated by chronic activation of DMH GABAergic neurons, suggesting that inhibitory input from the DMH can counteract stress-induced hyperactivity. Moreover, a negative correlation between firing rate and the coefficient of variation (CV) was observed across all groups (Fig. 3J, 3M and 3P). CUS also depolarized the resting membrane potential of POMC neurons in both sexes, this effect was attenuated by DMH GABAergic activation during chronic stress (Fig. 3Q).

## DISCUSSION

Despite extensive studies on POMC and AgRP neurons in response to hunger and satiety signals and their well-established roles in the regulation of food intake, their involvement in psychological stress responses, particularly in chronic stress–induced neuroendocrine and behavioral adaptations and maladapatations, remains poorly understood. POMC neurons in the ARC receive both GABAergic and glutamatergic inputs from local circuits as well as from other brain regions [1, 12, 15, 16]. Analysis of excitatory and inhibitory synaptic input onto POMC neurons revealed that chronic stress did not affect excitatory inputs but reduced inhibitory synaptic transmission [6]. This reduction was abolished when action potential–dependent transmission was blocked, indicating that CUS likely induces hyperpolarization of presynaptic GABAergic neurons or their terminals projecting to POMC neurons.

Our earlier studies showed that AgRP neurons are hyperpolarized by CUS [5]. Because AgRP neurons provide local GABAergic input to POMC neurons in ARC [10, 11, 14, 23], it was reasonable to expect that reduced inhibitory tone onto POMC neurons might result from decreased AgRP neuron activity. Surprisingly, chronic chemogenetic activation of AgRP neurons failed to attenuate stress-induced hyperactivity of POMC neurons. This result appears inconsistent with prior studies showing that acute optogenetic stimulation evokes GABAergic IPSCs in POMC neurons and robustly inhibits their firing [10, 14]. In our study, AgRP neurons were activated in vivo via CNO administered in drinking water throughout the 10-day stress exposure period, and POMC neurons were recorded 24 hours after the final stressor/CNO treatment. This design aimed to test whether sustained activation of AgRP neurons could mitigate stress-induced POMC hyperactivity. In contrast, optogenetic stimulation in vitro delivers acute, high-frequency, synchronized activation that efficiently drives GABA release and transiently suppresses POMC neuron firing. Several additional factors may explain the lack of effect of chronic chemogenetic activation. First, stress-induced hyperexcitability may be restricted to subpopulations of POMC neurons not innervated by AgRP terminals, limiting the efficacy of AgRP activation. Second, chronic stress could weaken synaptic efficacy at AgRP→POMC connections, thereby limiting inhibitory signaling. Finally, stress may engage additional upstream or parallel circuits that further diminish the functional impact of AgRP-derived inhibition. Together, these observations suggest that activation of AgRP neurons during chronic stress is insufficient to suppress the development of POMC neuron hyperactivity.

DMH contains a heterogeneous and intermingled neuronal population, including both glutamatergic and GABAergic neurons, with GABAergic neurons predominating [17, 18]. Evidence supports a critical role for the DMH in regulating stress responses. Previous studies in rats have shown that bilateral lesions of the DMH or blockade of glutamatergic transmission disinhibit neuroendocrine responses to emotional stressors [26–28], suggesting that the DMH normally exerts a inhibitory influence on neuroendocrine stress responses. One would therefore expect that chronic stress could alter the activity of DMH neurons, potentially reducing this inhibitory control and contributing to heightened stress reactivity. Using whole-cell patch-clamp recordings from GABAergic neurons in the DMH, we assessed the impact of chronic stress on neuronal activity and found that CUS suppresses firing in both male and female mice. Notably, while male and female mice displayed comparable populations of spontaneously active DMH GABAergic neurons, female neurons exhibited higher baseline firing rates and were more susceptible to stress-induced suppression than those in males. Sex differences in DMH GABAergic neurons extend beyond the magnitude of stress-induced suppression of firing rates, manifesting as distinct action potential properties in response to chronic stress. In males, CUS increased AP threshold and afterhyperpolarization amplitude, which would reduce neuronal excitability by requiring greater depolarization to initiate spikes and enhancing post-spike hyperpolarization. In contrast, in females, CUS increased AP half-width, rise time, and decay time, suggesting alterations in ion channel kinetics that may slow spike generation and propagation. These distinct electrophysiological adaptations reveal that chronic stress engages different cellular mechanisms in males and females to suppress DMH GABAergic neuron firing. Collectively, these findings reveal a previously unrecognized sex difference in the stress-induced suppression of DMH GABAergic neurons, evident in changes to their membrane properties and action potential dynamics. Given that DMH GABAergic neurons project to and inhibit ARC POMC neurons [15, 16], this suppression likely contributes to the reduced inhibitory tone and resulting hyperactivity of POMC neurons under chronic stress. This idea is supported by our finding that chronic chemogenetic activation of DMH GABAergic neurons during stress exposure effectively attenuates stress-induced POMC neuron hyperactivity in both sexes.

Interestingly, consistent with our observations that female DMH GABAergic neurons exhibit higher basal firing rates and are more strongly suppressed by chronic stress, we found that the population of spontaneously active POMC neurons was smaller in females than in males under basal conditions (62.5% vs. 76.9%) but increased more than in males following chronic stress (85.7% vs. 80.0%). These findings suggest that the higher basal activity of DMH GABAergic neurons in females tightly constrains POMC neuron activity under normal conditions, while their greater vulnerability to stress renders POMC neurons more susceptible to disinhibition when upstream inhibition is reduced. This stress-induced disinhibition of POMC neurons may, in turn, amplify their output, potentially altering downstream neuroendocrine and behavioral responses to stress.

Unlike AgRP neurons, which are almost exclusively GABAergic [23], POMC neurons in the ARC are heterogeneous, comprising glutamatergic and GABAergic subpopulations, as well as neurons that co-express both or neither neurotransmitter [29–34]. This diversity suggests that POMC neurons may play multiple, and potentially opposing, roles in regulating stress responses and other hypothalamic functions. It remains to be determined which subpopulations are selectively sensitive to emotional stress, whether they are preferentially innervated by DMH GABAergic neurons, and how their neurotransmitter identity shapes functional responses to chronic stress.

In summary, this study identifies DMH GABAergic neurons as a stress-sensitive inhibitory control node that constrains POMC neuron activity. Chronic stress suppresses DMH GABAergic firing, contributing to disinhibition and hyperactivity of POMC neurons, whereas chronic activation of DMH GABAergic neurons can counteract these stress-induced changes. This work uncovers a novel mechanism by which the DMH-ARC inhibitory circuit is reshaped under chronic stress. Importantly, the observed sex differences emphasize the complexity of this circuit-level regulation and may underlie sex-specific vulnerability to stress-related behavioral and neuroendocrine outcomes.

## REFERENCES

1. Bagnol, D., et al., Anatomy of an endogenous antagonist: relationship between Agouti-related protein and proopiomelanocortin in brain. J Neurosci, 1999. 19(18): p. RC26.

2. Lu, X.Y., et al., Diurnal rhythm of agouti-related protein and its relation to corticosterone and food intake. Endocrinology, 2002. 143(10): p. 3905–15.

3. Liu, J., et al., The melanocortinergic pathway is rapidly recruited by emotional stress and contributes to stress-induced anorexia and anxiety-like behavior. Endocrinology, 2007. 148(11): p. 5531–40.

4. Liu, J., et al., Melanocortin-4 receptor in the medial amygdala regulates emotional stress-induced anxiety-like behaviour, anorexia and corticosterone secretion. Int J Neuropsychopharmacol, 2013. 16(1): p. 105–20.

5. Fang, X., et al., Chronic unpredictable stress induces depression-related behaviors by suppressing AgRP neuron activity. Mol Psychiatry, 2021. 26(6): p. 2299–2315.

6. Fang, X., et al., Increased intrinsic and synaptic excitability of hypothalamic POMC neurons underlies chronic stress-induced behavioral deficits. Mol Psychiatry, 2023. 28(3): p. 1365–1382.

7. Qu, N., et al., A POMC-originated circuit regulates stress-induced hypophagia, depression, and anhedonia. Mol Psychiatry, 2020. 25(5): p. 1006–1021.

8. Lei, Y., et al., SIRT1 in forebrain excitatory neurons produces sexually dimorphic effects on depression-related behaviors and modulates neuronal excitability and synaptic transmission in the medial prefrontal cortex. Mol Psychiatry, 2020. 25(5): p. 1094–1111.

9. Lei, Y., et al., Leptin enhances social motivation and reverses chronic unpredictable stress-induced social anhedonia during adolescence. Mol Psychiatry, 2022. 27(12): p. 4948–4958.

10. Atasoy, D., et al., Deconstruction of a neural circuit for hunger. Nature, 2012. 488(7410): p. 172–7.

11. Dicken, M.S., A.R. Hughes, and S.T. Hentges, Gad1 mRNA as a reliable indicator of altered GABA release from orexigenic neurons in the hypothalamus. Eur J Neurosci, 2015. 42(9): p. 2644–53.

12. Cowley, M.A., et al., Leptin activates anorexigenic POMC neurons through a neural network in the arcuate nucleus. Nature, 2001. 411(6836): p. 480–4.

13. Wu, Q., et al., Starvation after AgRP neuron ablation is independent of melanocortin signaling. Proc Natl Acad Sci U S A, 2008. 105(7): p. 2687–92.

14. Rau, A.R. and S.T. Hentges, The Relevance of AgRP Neuron-Derived GABA Inputs to POMC Neurons Differs for Spontaneous and Evoked Release. J Neurosci, 2017. 37(31): p. 7362–7372.

15. Rau, A.R. and S.T. Hentges, GABAergic Inputs to POMC Neurons Originating from the Dorsomedial Hypothalamus Are Regulated by Energy State. J Neurosci, 2019. 39(33): p. 6449–6459.

16. Wang, D., et al., Whole-brain mapping of the direct inputs and axonal projections of POMC and AgRP neurons. Front Neuroanat, 2015. 9: p. 40.

17. Vong, L., et al., Leptin action on GABAergic neurons prevents obesity and reduces inhibitory tone to POMC neurons. Neuron, 2011. 71(1): p. 142–54.

18. Cullinan, W.E., D.R. Ziegler, and J.P. Herman, Functional role of local GABAergic influences on the HPA axis. Brain Struct Funct, 2008. 213(1-2): p. 63–72.

19. Cullinan, W.E., et al., Pattern and time course of immediate early gene expression in rat brain following acute stress. Neuroscience, 1995. 64(2): p. 477–505.

20. Canteras, N.S., et al., Severe reduction of rat defensive behavior to a predator by discrete hypothalamic chemical lesions. Brain Res Bull, 1997. 44(3): p. 297–305.

21. Nakamura, K., Y. Nakamura, and N. Kataoka, A hypothalamomedullary network for physiological responses to environmental stresses. Nat Rev Neurosci, 2022. 23(1): p. 35–52.

22. Flak, J.N., et al., Identification of chronic stress-activated regions reveals a potential recruited circuit in rat brain. Eur J Neurosci, 2012. 36(4): p. 2547–55.

23. Tong, Q., et al., Synaptic release of GABA by AgRP neurons is required for normal regulation of energy balance. Nat Neurosci, 2008. 11(9): p. 998–1000.

24. Lei, Y., et al., Neuronal HDAC9: A key regulator of cognitive and synaptic aging, rescuing Alzheimer’s disease-related phenotypes. bioRxiv, 2025.

25. Lei, Y., et al., Chronic social isolation-unpredictable stress induces early-onset cognitive deficits and exacerbates Abeta accumulation in the 5xFAD mouse model of Alzheimer’s disease. Mol Psychiatry, 2025. 30(10): p. 4720–4735.

26. Ebner, K., P. Muigg, and N. Singewald, Inhibitory function of the dorsomedial hypothalamic nucleus on the hypothalamic-pituitary-adrenal axis response to an emotional stressor but not immune challenge. J Neuroendocrinol, 2013. 25(1): p. 48–55.

27. Herman, J.P., et al., Central mechanisms of stress integration: hierarchical circuitry controlling hypothalamo-pituitary-adrenocortical responsiveness. Front Neuroendocrinol, 2003. 24(3): p. 151–80.

28. Boudaba, C., K. Szabo, and J.G. Tasker, Physiological mapping of local inhibitory inputs to the hypothalamic paraventricular nucleus. J Neurosci, 1996. 16(22): p. 7151–60.

29. Dicken, M.S., R.E. Tooker, and S.T. Hentges, Regulation of GABA and glutamate release from proopiomelanocortin neuron terminals in intact hypothalamic networks. J Neurosci, 2012. 32(12): p. 4042–8.

30. Jones, G.L., et al., Selective Restoration of Pomc Expression in Glutamatergic POMC Neurons: Evidence for a Dynamic Hypothalamic Neurotransmitter Network. eNeuro, 2019. 6(2).

31. Hentges, S.T., et al., GABA release from proopiomelanocortin neurons. J Neurosci, 2004. 24(7): p. 1578–83.

32. Hentges, S.T., et al., Proopiomelanocortin expression in both GABA and glutamate neurons. J Neurosci, 2009. 29(43): p. 13684–90.

33. Mazier, W., et al., mTORC1 and CB1 receptor signaling regulate excitatory glutamatergic inputs onto the hypothalamic paraventricular nucleus in response to energy availability. Mol Metab, 2019. 28: p. 151–159.

34. Wittmann, G., E. Hrabovszky, and R.M. Lechan, Distinct glutamatergic and GABAergic subsets of hypothalamic pro-opiomelanocortin neurons revealed by in situ hybridization in male rats and mice. J Comp Neurol, 2013. 521(14): p. 3287–302.

